# DeepArk: modeling *cis*-regulatory codes of model species with deep learning

**DOI:** 10.1101/2020.04.23.058040

**Authors:** Evan M. Cofer, João Raimundo, Alicja Tadych, Yuji Yamazaki, Aaron K. Wong, Chandra L. Theesfeld, Michael S. Levine, Olga G. Troyanskaya

## Abstract

To enable large-scale analyses of regulatory logic in model species, we developed DeepArk (https://DeepArk.princeton.edu), a set of deep learning models of the *cis*-regulatory codes of four widely-studied species: *Caenorhabditis elegans, Danio rerio*, *Drosophila melanogaster*, and *Mus musculus*. DeepArk accurately predicts the presence of thousands of different context-specific regulatory features, including chromatin states, histone marks, and transcription factors. In vivo studies show that DeepArk can predict the regulatory impact of any genomic variant (including rare or not previously observed), and enables the regulatory annotation of understudied model species.

## Main

Deciphering the regulatory function of the non-coding genome remains a grand challenge of modern biology. Model species have long been at the forefront of biological discovery and biomedical innovation^1^, but our knowledge of the *cis*-regulatory logic remains incomplete. Many important questions remain: how should we mutate a fly enhancer to change its activity in a tissue-specific manner? Which regulatory variants for mouse disease genes are functional? How can we predictively edit the genome to efficiently guide experimentation? Answering these questions requires interpreting specific effects of any genomic variant, including changes to chromatin states, histone modifications, and binding of transcription factors. Addressing this challenge across the entire spectrum of genomic variation requires generalizing from the experimental studies (e.g. ChIP-Seq data) to learn the regulatory code and thus enable the prediction of effects of any genomic variant. These effects must be predicted in specific contexts including developmental stage, cell and tissue type, and drug treatments - an experimentally intractable set of combinations.

Existing approaches for model organisms fall short of this goal. A common approach is to scan for highly conserved binding sites with position weight matrices. However, such motifs have limited context information and fail to consider the multiple interacting factors that frequently delineate histone marks or chromatin accessibility^2,3^. In contrast, sequence-based deep learning models are capable of learning this context-specific *cis*-regulatory code, and while they have proven to be powerful for the complex task of predicting regulatory activity from genomic sequences^3–5^, their applicability to model organisms remains largely undemonstrated aside from a few limited exceptions^6,7^.

We developed a set of sequence-based deep convolutional neural networks (CNNs), which we collectively named “DeepArk”, modeling the *cis*-regulatory codes of four of the most widely-studied model organisms: *Caenorhabditis elegans*, *Danio rerio, Drosophila melanogaster*, and *Mus musculus* (**Fig. 1a**). To the best of our knowledge, DeepArk is the first such resource for these model organisms. Intuitively, DeepArk provides an *in silico* ChIP-seq capability: given a genomic sequence as input, DeepArk’s CNNs predict the activity of a total of 6,562 regulatory features, including histone marks, different transcription factors (TFs), RNA polymerases, and chromatin accessibility (**Supplementary Table 1**). Notably, DeepArk leverages a wide sequence context of 4,095 bp to provide accurate predictions for broad regulatory features with complex regulatory origins (e.g. chromatin accessibility). DeepArk’s multitask approach to modeling also allows it to make such predictions efficiently. Many predictions are made in specific contexts - larval or adult stages, specific tissues or cell types, and under particular treatments (e.g. lipopolysaccharide stimulation). Importantly, for most of the organisms and regulatory features considered, DeepArk is the first method capable of accurately predicting regulatory activity from genomic sequence and the regulatory effects of genomic variants^6,7^.

**Figure 1:**
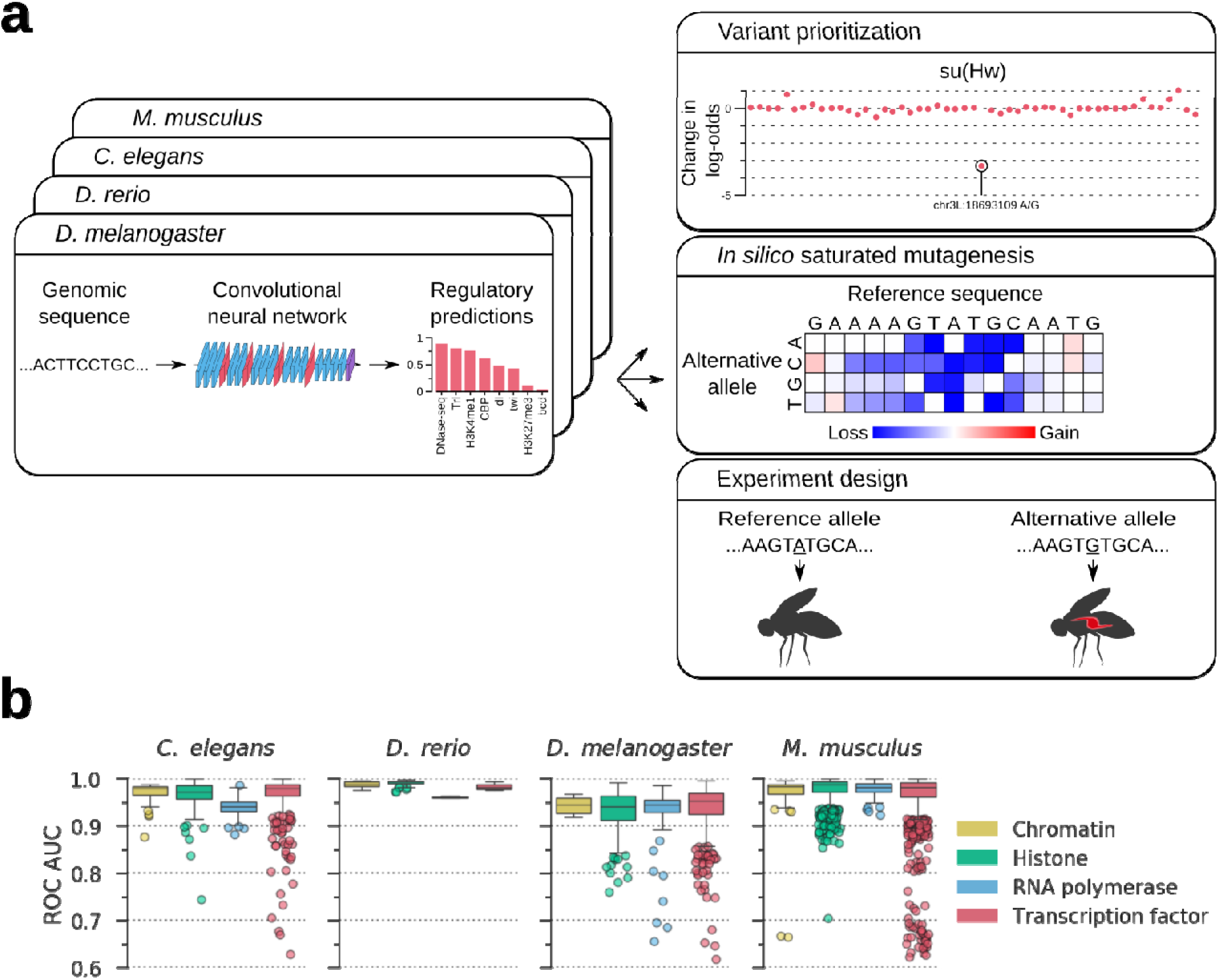
Overview of DeepArk models and their predictive accuracy. **(a)** The DeepArk architecture (**Supplementary Fig. 1**) uses convolutional layers to scan an input sequence for regulatory motifs, and maximum pooling layers to perform dimensionality reduction. By utilizing many successive layers, DeepArk is able to extract complex motifs, interactions between motifs, and consider a wide sequence context of 4,095 bp. Key applications enabled by DeepArk include prioritizing observed genomic variants by their putative regulatory effects (top right), exposing the predictive sequence features for regulatory events through *in silico* saturated mutagenesis (middle right), and predicting the regulatory effects of novel variants for prospective experiments (bottom right). **(b)** Performance on test chromosomes from each organism, as quantified by the area under the curve (AUC) of the receiver operating characteristic (ROC) curve. Only regulatory features with at least 50 positive test examples are included. For each box plot, the center line marks the median, and the top and bottom edges of the box mark the first and third quartiles respectively. The top and bottom whiskers extend to 1.5× the interquartile range (IQR), with data points outside of this range considered outliers and plotted individually.

We trained each DeepArk model on publicly-available genome-wide measurements of regulatory activity (i.e. ChIP-seq of TFs and histone marks, DNase-seq, and ATAC-seq) from its respective species, and tested its performance on chromosomes that were not used during training (Online Methods and **Supplementary Table 2**). Consistent with the accurate inference of *cis*-regulatory logic, we found that each DeepArk model correctly predicted the regulatory activity of the test sequences (**Fig. 1b** and **Supplementary Table 1**).

At its core, DeepArk provides a mapping from DNA sequences to regulatory activities. By comparing DeepArk’s predictions for separate sequences, we can identify how sequence differences lead to regulatory differences. As a consequence, we can predict whether a variant might increase or decrease regulatory activity for any of DeepArk’s 6,562 regulatory features. This ability to predict the *cis*-regulatory effects of genomic variants is an important step forward for model species genomics, as there is a paucity of such methods available.

To assess DeepArk’s ability to guide the interpretation of regulatory variants, we compared its predictions for the regulatory effects of variants in an enhancer of *ALDOB* with actual effects as measured by Patwardhan et al. in a massively parallel *in vivo* reporter assay (MPRA) in murine livers^8^. By barcoding each variant and quantifying enhancer activity with RNA-sequencing, the MPRA can test the expression-modulating effects of all possible single nucleotide polymorphisms (SNPs) in a given enhancer. DeepArk’s mouse model’s variant effect predictions were significantly correlated with the expression effects of the SNPs measured in the *ALDOB* enhancer MPRA (Pearson’s *r*=0.714, P=3.58×10^−122^ and Spearman’s *ρ*=0.587, P=2.91 ×10^−73^, n=777) (**Supplementary Fig. 2** and **Supplementary Table 3**), further demonstrating that DeepArk’s predictions accurately reflect *in vivo* observations.

DeepArk can also be deployed to investigate regulatory loci at the genome- or chromosome-wide scale. For example, a researcher interested in identifying loci guiding the spreading of the dosage compensation complex (DCC) of *C. elegans*, a complex that both binds and spreads along the inactivated X chromosome^9^. They could use DeepArk to investigate the DCC computationally and identify sites involved in the DCC’s initial recruitment. First, the researcher identifies a region as a highly-probable site of DCC binding by scanning all of chromosome X for binding of several protein components of the DCC (e.g. *dpy-27*) *in vivo* (**Supplementary Table 4** and **Supplementary Fig. 3**). Conducting an *in silico* saturated mutagenesis of the putative DCC-bound region for DCC members reveals a single highly constrained sequence (GCGCAGGGA) that is necessary for DCC binding *in vivo* (**Supplementary Fig. 4**) and consistent with existing literature^10^. Thus, DeepArk may be used to interpret the binding patterns of even relatively complicated protein complexes.

As another application, DeepArk can directly assist in studying regulatory genomics. We used the DeepArk model for *D. melanogaster* to investigate the regulatory effects of mutations in the mesodermal enhancer of the *T48* gene^11^ whose timely expression regulates gastrulation in flies^11,12^ (**Supplementary Fig. 5** and **Supplementary Tables 5** and **6**) and relies on binding of zelda, a pioneer factor^13,14^. DeepArk predicted that the original suboptimal zelda binding site would have the lowest probability for zelda binding, whilst the variants CTT>CTA and CTT>GTA would have moderate probability and the CTT>CTG variant, the largest positive effect on zelda binding. To test the predictive capabilities of DeepArk, we examined the *in vivo* expression in live embryos of these three variants. Experimental quantification of the total transcriptional output of the T48 enhancer variants clearly shows that DeepArk’s predictions were accurate, and that, as expected, an increase in zelda binding at probability correlates with an increase in gene activation *in vivo* (**Fig. 2**). This both experimentally confirmed DeepArk’s predictions and demonstrated its utility in designing genome editing experiments.

**Figure 2:**
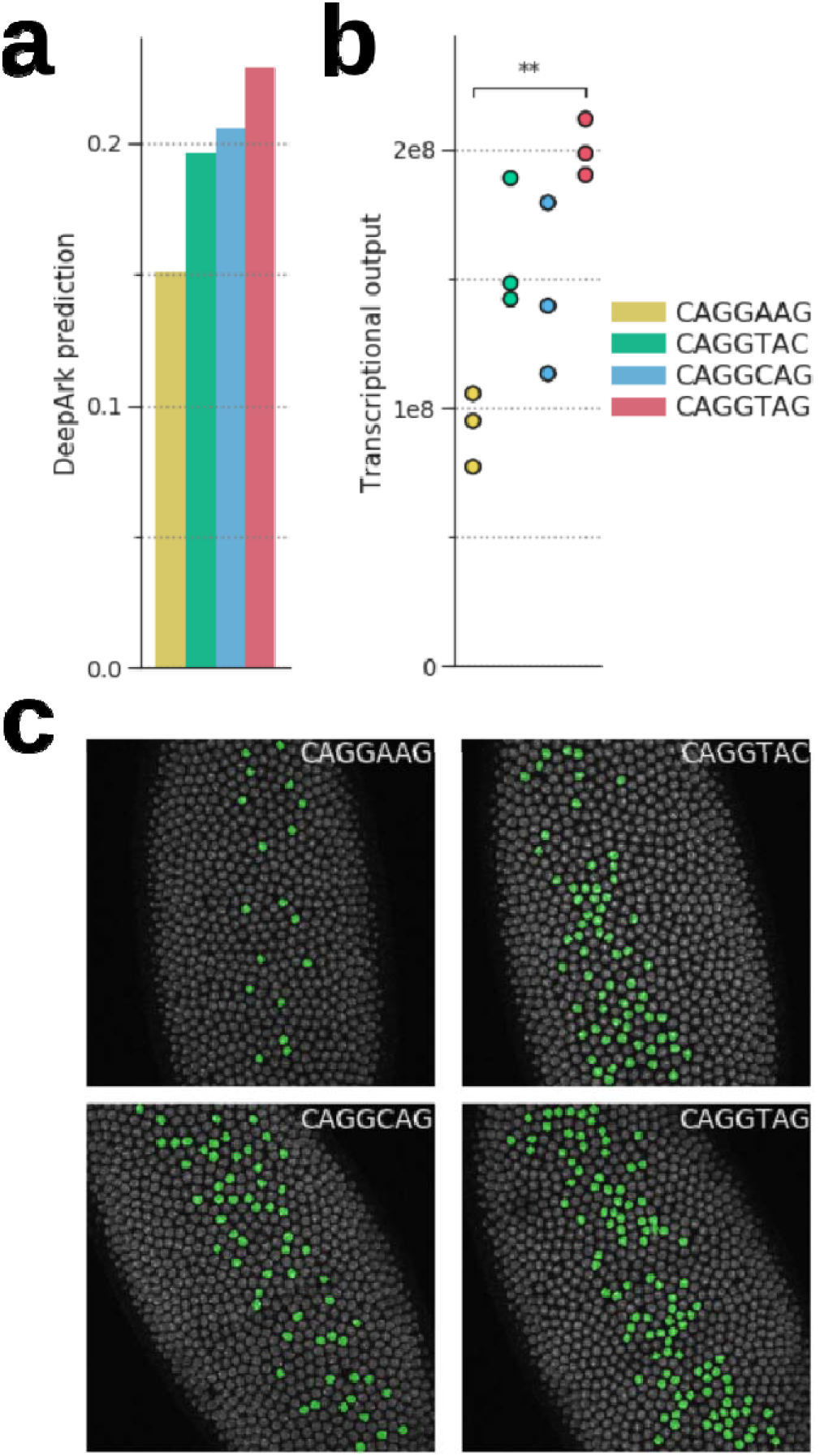
DeepArk’s predicted effects for the different T48 mesodermal enhancer variants correlate with *in vivo* results. **(a)** Plot of DeepArk predictions for Zelda binding during nuclear cycle 14 (accession no. SRX858993) for each of the four enhancer alleles. The CAGGTAG allele has the highest predicted probability of binding, with the reference allele CAGGAAG exhibiting the lowest. **(b)** The total transcriptional output for each of the four alleles, as quantified with *in vivo* MS2 tagging during nuclear cycle 14. Note that CAGGTAG and CAGGAAG have the lowest and highest transcriptional outputs respectively, which is consistent with DeepArk’s predictions. Bonferroni-corrected two-sided t-test with unequal variances, **P = 4.139 × 10^−3^; all others, P >5 × 10^−2^. **(c)** False colored nuclei with active transcription in *Drosophila* embryos during minute 20 of nuclear cycle 14 illustrate the distinct levels of transcriptional activation induced by each allele.

DeepArk may also be particularly useful for researchers of understudied model organisms without available regulatory data. Presently, pairwise alignments of regulatory regions allow the detection of constrained non-coding sequences in the absence of regulatory assays, but they are only a proxy for regulatory function and their interpretation can be confounded. For instance, some regulatory elements are enriched for recent evolution^15^, while other highly conserved non-coding regions have no known function^16^. By directly predicting regulatory activity from sequence, DeepArk alleviates this challenge. To that end, we used the DeepArk model for the model organism *D. rerio* to predict chromatin accessibility and H3K4me3 marks during development in the genome of *Oryzias latipes*, or Medaka, a fish that diverged from *D. rerio* an estimated 314 to 332 million years ago^17^. Even after filtering conserved loci, we find that DeepArk accurately predicts ChIP-seq and ATAC-seq peaks for developing *O. latipes* (average ROC AUC of 0.927) using only their genomic sequence as model input (**Supplementary Table 7**). Thus DeepArk may also be used to help annotate the genomes of understudied organisms when whole-genome assays of regulatory features do not already exist (**Supplementary Fig. 6**).

As we have shown, DeepArk excels at a number of diverse tasks such as accurately predicting the regulatory landscape of model species and predictive genome editing. Here we have shown a few examples of DeepArk’s utility, which we have made publicly available (https://DeepArk.princeton.edu) to enable new and more efficient approaches for experimental studies. Since we developed DeepArk in a transparent and open-source manner, we anticipate that it may be extended and repurposed for other tasks via transfer learning. Furthermore, DeepArk could be used as a scoring function in a sequence optimization and design pipeline^18^, as covariates for models of more complicated regulatory events such as enhancer-promoter looping^19^, and in high-resolution association mapping of animal models as it becomes widespread^20^. Thus, DeepArk will contribute to a number of diverse experimental and computational analyses, both directly through its predictions or as part of larger computational pipelines.

## Online Methods

### DeepArk model architecture

DeepArk is a collection of four deep convolutional neural networks, each modeling the activity of different regulatory features in a separate model organism. In total, DeepArk is capable of making predictions for 6,562 context-specific regulatory features. In what follows, we detail the design and structure of the DeepArk model.

Each DeepArk model takes a 4095 bp genomic sequence as input, and predicts the probability that the centermost base of this sequence is covered by a peak for each regulatory feature of interest. This input sequence is encoded as a 4095×4 one-hot matrix with columns corresponding to each base in the sequence, and rows corresponding to adenine, cytosine, guanine, and thymine respectively. The output of each DeepArk model is a vector of length *N*, where *N* is the number of features for that model’s given organism (**Supplementary Table 1**). DeepArk is a multitask model, which means it jointly learns the sequence-specific activities of multiple regulatory features simultaneously. The DeepArk architecture was fixed across organisms (**Supplementary Fig. 1**), but we learned distinct model parameters and hyperparameters for each organism (**Supplementary Table 8**).

The DeepArk architecture (**Supplementary Fig. 1**) consists of a deep convolutional neural network, wherein the network’s output is the functional composition of many linear and non-linear transformations, called “layers”. The specific parameters of these transformations are selected during training to optimize the objective function. We consider four types of transformations in our network: convolutional layers, maximum pooling layers, batch normalization layers, and the rectified linear unit (ReLU) and sigmoid activation functions. The basic unit of our model is a multi-layer “convolutional unit”, which contains, in order, a batch normalization layer, a ReLU layer, and a convolution layer. We further organize our model into five multi-layered convolutional blocks. We used maximum pooling at the start of each convolutional block, as we found that spatial invariance and reduced training time allowed us to improve our model. The output of the final convolutional block is fed into a length-1 convolution with output channels equal to the number output features of the model, and fed into the sigmoid activation function.

We designed the DeepArk architecture to regularize it and avoid overfitting. First, we averaged predictions made for the forward and reverse complement of sequences. Second, we leveraged spatial dropout^21^, which randomly zeros out channels in the input to a convolutional layer. Typically, dropout^22^ randomly zeros sets input values to zero with probability *P*, which has the effect of forcing the model to overcome perturbations in internal values (i.e. without altering the sequence input) to make correct predictions. However, highly correlated sequence positions in convolutional neural network inputs may diminish the effectiveness of dropout and slow training. Conversely, spatial dropout mitigates this by zeroing out entire channels of the convolutional layer’s input.

### Training examples

Training examples are 2-tuples of a 4095 bp genomic sequence and a label vector. For each example, a given feature’s entry in the label vector is positive if the center base of the 4095 bp sequence is overlapped by a peak from the feature’s corresponding ChIP-seq, DNase-seq, or ATAC-seq experiment. With the exception of ENCODE blacklisted regions^23^, all positions in the genome were considered valid examples.

Non-intersecting training, validation, and testing sets were generated by whole-chromosome holdout (**Supplementary Table 2**). Validation data were generated by randomly drawing 64,000 examples from a given species’s set of validation chromosomes. Training and validation examples are drawn uniformly and with replacement. Each species’ test set consisted of 1 million examples drawn uniformly and without replacement from the held-out test chromosomes for said species. Only features with at least 50 positive examples in the held-out test data were considered when calculating performance metrics.

### Training DeepArk

We used stochastic gradient descent with momentum and mini-batches of 128 examples to select network weights that optimized the model objective function during training. Specifically, our objective function was the sum of the binary cross-entropy (BCE) loss and an L2 regularization term,

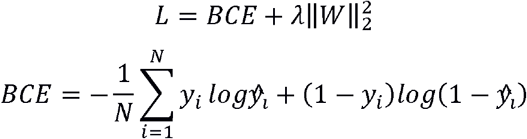

where *y_i_* is the vector of target labels for example *i*, 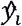 is DeepArk’s prediction for example *i*, λ is the weight decay hyperparameter, *W* is the weight matrix, and *N* is the mini-batch size. Model validation performance was evaluated every 5000 training steps with a validation set drawn randomly from a set of held-out validation chromosomes (see “Training examples” section in Online Methods). When minimum validation loss failed to decrease for five consecutive epochs, we decrease the learning rate by 20% of its current value. We terminated training when validation loss stopped decreasing for a sustained period of time. Hyperparameters for SGD and model training (**Supplementary Table 8**) were selected based on each model’s performance on its respective validation set. DeepArk was implemented and trained with Selene^24^.

### Training data preparation

Labels for training, validation, and testing data were constructed using publicly-available ChIP-seq, DNase-seq, and ATAC-seq for *C. elegans*, *D. rerio*, *D. melanogaster*, and *M. musculus*. For *C. elegans*, *D. melanogaster*, and *M. musculus*, we used peak intervals from ChIP-atlas^25^ - a large compendium of uniformly processed high-throughput sequencing experiments sourced from the Sequence Read Archive (SRA), the European Nucleotide Archive (ENA), and the DNA Data Bank of Japan (DDBJ). Specifically, we used peaks called with a maximum Q-value cutoff of 1×10^−5^.

To keep our methodology consistent, we called our own peaks for *D. rerio*. Specifically, we downloaded aligned BAMs for ChIP-seq and ATAC-seq experiments from the DANIO-CODE website^26^. Peaks were called for these BAMs using MACS2^27^ with a maximum Q-value cutoff of 1×10^−5^ and an effective genome size of 8.1×10^8^. This approximate effective genome size was calculated by counting the number of unambiguous bases in the Genome Reference Consortium Zebrafish Build 11 (GRCz11), without including repeats^28^. The repeat-masked genome was downloaded from UCSC genome browser annotation database^29^.

To ensure that we only considered high-quality experiments, we removed those with too few peaks, an insufficient number of mapped reads, or average read length shorter than 32 bases pairs (**Supplementary Table 9**). We also removed experiments that did not list a specific antibody target. We manually curated sample metadata regarding strains, cell lines, genetic modifications, and sample treatment. Since there exists a wide range of mouse cell lines and strains with extensive genetic and phenotypic diversity among them, we removed mouse experiments that did not reference a specific strain or cell line.

Finally, we removed experiments where there was duplication or redundancy between SRA, ENA and DDBJ. We considered experiments to be duplicates if they were from the same species, differed by fewer than 100 peaks, and had the same number of unmapped, mapped, and duplicate-free reads. We manually inspected the FASTQ files to ensure true duplication. In cases where both accessions had the same metadata, we discarded one of the duplicate accessions at random. If the duplicates did not have the same antibody or biological source (e.g. cell type) listed, we discarded all of them.

### Analysis of massively parallel reporter assay

To demonstrate DeepArk’s accuracy, we used it to predict the regulatory effects of all possible variants in the ALDOB enhancer^8^. We downloaded variant effects for the massively parallel *in vivo* reporter assay of the ALDOB from MaveDB^30^. The predicted functional effect of variants was calculated with *in silico* saturated mutagenesis. Specifically, the change in chromatin accessibility (accession no. SRX3201109) at the center of the 259 bp ALDOB enhancer (hg19:chr9:104195570-104195828) was predicted for all possible variants within the 4,095 bp window, and those reported in the MPRA were retained for analysis.

### Predicting the binding of the DCC in *C. elegans*

To identify a high-confidence binding site for the DCC, we scanned the entire X chromosome and made predictions every 200 bp with DeepArk. We took the mean probability across all features corresponding to DCC components (**Supplementary Table 4**) as a proxy of DCC binding probability. The site with the maximum mean probability across DCC components was then analyzed with *in silico* saturated mutagenesis (**Supplementary Table 10**).

### Cloning of T48 enhancer MS2 reporter alleles

To clone the four different T48 enhancer MS2 reporters the T48 enhancer was first cut with NotI from the T48>MS2>yellow plasmid^11^ and subcloned into a pGEM-T Easy vector. Site-directed mutagenesis was then performed by amplifying the pGEM-T Easy T48 enhancer vector (**Supplementary Table 11**). The different PCR reactions were digested with Dpn and transformed into e.coli in order to obtain clones of the four different T48 alleles. These plasmids were then individually subcloned into the pbphi-evePr-MS2-yellow vector^31^ using NotI.

### Live imaging of T48 enhancer alleles

To visualize live transcription of the T48 MS2 reporters, female fly virgins carrying the nanos>MCP-GFP and His2Av-mRFP fusion proteins^32^ were crossed to males carrying the MS2 reporter genes inserted on a landing site of the third chromosome (strain 9450, Bloomington stock center). The resulting embryos were dechorionated and mounted between a semipermeable membrane and a coverslip with Halocarbon oil 27 (Sigma). Embryos were imaged from the beginning of nuclear cycle 14 up to the onset of gastrulation using a Zeiss LSM 880 confocal microscope and a Plan-Apochromat 40x/1.3 NA oil-immersion objective. For each time point a stack of 21 images separated by 0.5 μm with a final time resolution of 14 seconds was acquired at 16 bit. Two laser lines at 488nm and 561nm were used to excite the green and red fluorophores, respectively. The same imaging conditions were used across the three replicates and the four different reporter lines.

### Image analysis and transcription quantification

To quantify the fluorescent signal resulting from the embryos live transcription, the 21 images corresponding to each time point were converted into maximum projections. The subsequent analysis was processed by segmenting the nuclei using the His2Av-mRFP channel and tracking the segmented individual nuclei during nuclear cycle 14. To record the MS2-GFP fluorescent signal corresponding to the transcription foci, an average of the signal for the three brightest pixels within each nucleus was determined after subtracting the background GFP signal. Total transcriptional output was calculated by adding the transcription foci signal for each active nuclei for the first 200 time frames of each embryo after the onset of nuclear cycle 14. Please refer to Fukaya et al.^31^ for a more detailed description of the image analysis methods.

### Interspecies regulatory prediction

To illustrate DeepArk’s ability to make accurate predictions in novel species (i.e. not *C. elegans, D. rerio*, *D. melanogaster*, or *M. musculus*), we used the DeepArk model for *D. rerio* to predict regulatory activity of sequences from the genome of *O. latipes*, which diverged from *D. rerio* between 314 and 332 million years ago^17^. In specific, we used leveraged extant ATAC-seq and H3K4me1 ChIP-seq data from *O. latipes*.

To generate testing examples for *O. latipes*, we randomly drew 1 million locations from the *O. latipes* reference genome without replacement. We ignored regions that contained an excess (i.e. >50) of ambiguous bases. To ensure the model was truly generalizing to the *O. latipes* genome, we removed sequences that were conserved between *D. rerio* and *O. latipes*. To identify conserved bases, we used a multiple whole-genome alignment of eight vertebrates - including *O. latipes* - to the *D. rerio* reference genome. To enable comparisons between the two fish, morphological stages of *O. latipes* development were matched to their corresponding stages in *D. rerio^33^*.

Labels for the testing examples were assigned using existing ATAC-seq and ChIP-seq data from *O. latipes*. We downloaded unprocessed FASTQ files from SRA using the SRA toolkit^34^. We filtered and clipped reads using TrimGalore^35^. We used BWA-MEM^36^ to align reads to the *O. latipes* reference genome^17^. Following alignment, we used SAMtools^37^ to index and sort the BAM files, and the ‘MarkDuplicates’ command from Picard Tools^38^ to identify and remove duplicate reads in each BAM file. Finally, we used MACS2^27^ to call peaks with a Q-value cutoff 1×10^−5^ and an effective genome size of 8.18×10^8^.

Lastly, RNA-seq data for *D. rerio* and *O. latipes* were used to visualize changes in expression and compare to changes in histone modifications at promoters (**Supplementary Fig. 6**). We downloaded the unprocessed FASTQ files for these data from SRA using the SRA toolkit^34^. Using HISAT2^39^, the processed reads for *O. latipes* were aligned to its reference genome^17^, and the reads for *D. rerio* were aligned to GRCz11^28^. Coverage was quantified as counts per million mapped reads (CPM) using the ‘bamCoverage’ command from deepTools^40^.

## Code availability

DeepArk is freely accessible through our user-friendly web server (https://DeepArk.princeton.edu). The code to run DeepArk locally is also available on GitHub (https://github.com/FunctionLab/DeepArk).

## Data availability

All ChIP-atlas data used for training DeepArk models are from the ChIP-atlas website (https://chip-atlas.org/). Training data for *C. elegans, D. melanogaster*, and *M. musculus* were downloaded from the following URLs:

- http://dbarchive.biosciencedbc.jp/kyushu-u/ce10/allPeaks_light/allPeaks_light.ce10.05.bed.gz
- http://dbarchive.biosciencedbc.jp/kyushu-u/dm3/allPeaks_light/allPeaks_light.dm3.05.bed.gz
- http://dbarchive.biosciencedbc.jp/kyushu-u/mm9/allPeaks_light/allPeaks_light.mm9.05.bed.gz

All training data for *D. rerio* are from the DANIO-CODE website (https://danio-code.zfin.org). Their URLs are listed in Supplementary Table 12. All data for *Oryzias latipes* are from SRA. Their accessions are listed in Supplementary Table 7. Murine MPRA data are from MaveDB (https://www.mavedb.org/scoreset/urn:mavedb:00000006-a-1/). The 8-way multiple genome alignment used to compare the genomes of *D. rerio* and *O. latipes* is from the UCSC Genome Browser website (https://hgdownload.soe.ucsc.edu/goldenPath/danRer7/multiz8way/multiz8way.maf.gz). Developmental RNA-seq data for *D. rerio* (accession no. SRX3353221) and *O. latipes* (accession no. SRX3353227) used in Supplementary Figure 6 were downloaded from SRA. Raw videos from imaging are available on Zenodo (https://doi.org/10.5281/zenodo.3759736). Predictions for DCC component binding along the *C. elegans* X chromosome are also available on Zenodo (https://doi.org/10.5281/zenodo.3759699).

## Acknowledgements

The authors acknowledge all members of the Troyanskaya and Levine labs for their helpful discussions. They would also like to thank Beryl M. Jones, Siena Dumas Ang, and Rachel Kaletsky for their feedback regarding the DeepArk web server. The authors are pleased to acknowledge that this work was performed using the high-performance computing resources at Simons Foundation and the TIGRESS computer center at Princeton University. E.M.C. was supported by National Institutes of Health (NIH) grant T32 HG003284 and the National Science Foundation Graduate Research Fellowship Program (NSF-GRFP). This work was supported by NIH grant R01 GM071966 (O.G.T.) and R35 GM118147 (M.S.L.). O.G.T. is a senior fellow of the Canadian Institute for Advanced Research (CIFAR) Genetic Networks program.

## Author contributions

E.M.C. conceived the idea; designed, implemented, and developed the DeepArk models; and performed all analyses. J.R. designed and executed all imaging experiments. Y.Y. designed and prepared the T48 reporters. E.M.C., A.T., and A.K.W. designed, implemented, and deployed the DeepArk web server. E.M.C., C.L.T., J.R., M.S.L., and O.G.T. wrote the manuscript with feedback from all other authors.

## Competing interests

The authors declare that no competing interests, financial or otherwise, exist.

## Notes

https://DeepArk.princeton.edu

### Competing Interest Statement

The authors have declared no competing interest.

### Summary of Updates

The missing supplemental tables have been added.

