## Supplementary Material for "DeepArk: modeling *cis*-regulatory codes of model species with deep learning"

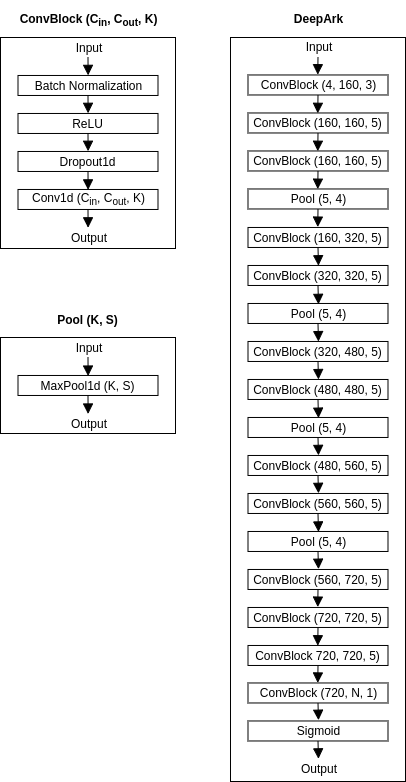

**Supplementary Figure 1: Overview of DeepArk model architecture.** The architecture used for DeepArk. Convolutional blocks have *C_in_* input channels, *C_out_* output channels, and a kernel size of *K*, while pooling blocks have a kernel size of *K* and stride of *S.* The dropout rate used by spatial dropout layers during training varied by species (**Supplementary Table 8**).

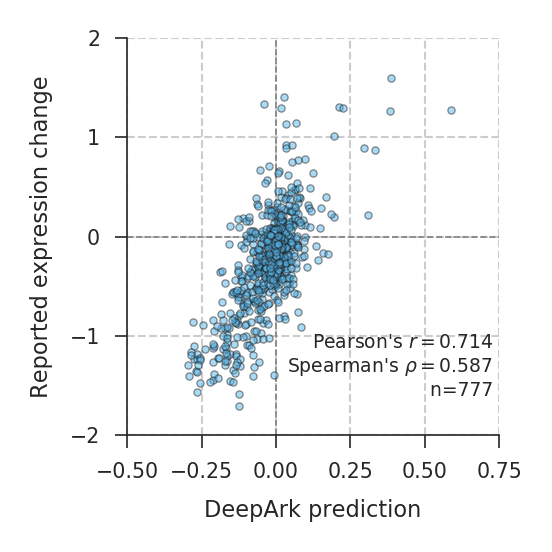

**DeepArk’s predictions significantly correlate with variant effects measured in a massively parallel reporter assay of enhancer activity.** Shown are DeepArk’s predictions for all possible variants in the *ALDOB* enhancer (hg19:chr9:104195570-104195828) and the expression effects recorded from the MPRA from Patwardhan et al.^8^ (**Supplementary Table 3**). Correlation between the two values is strong (Pearson’s r=0.714, P=3.58×10-122, n=777 and Spearman’s ρ=0.587, P=2.91×10-73, n=777). DeepArk’s predictions are for DNase-seq from the liver of an untreated male CD-1 mouse (accession no. SRX3201109).

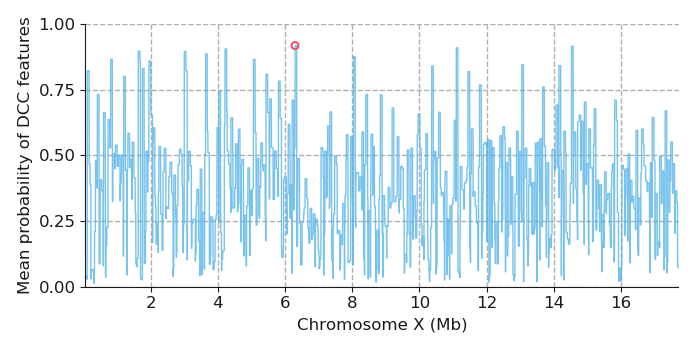
**Supplementary Figure 3:** **DCC-bound regions of the *C. elegans* X chromosome predicted by DeepArk.** The mean predicted probability of DCC features (**Supplementary Table 4**) throughout the X chromosome of *C. elegans*. To enhance readability, we plot the maximum value per 50 Kb bin of the chromosome. The position with the maximum predicted probability (ce11:chrX:6294496-6298590) is marked with a red circle.

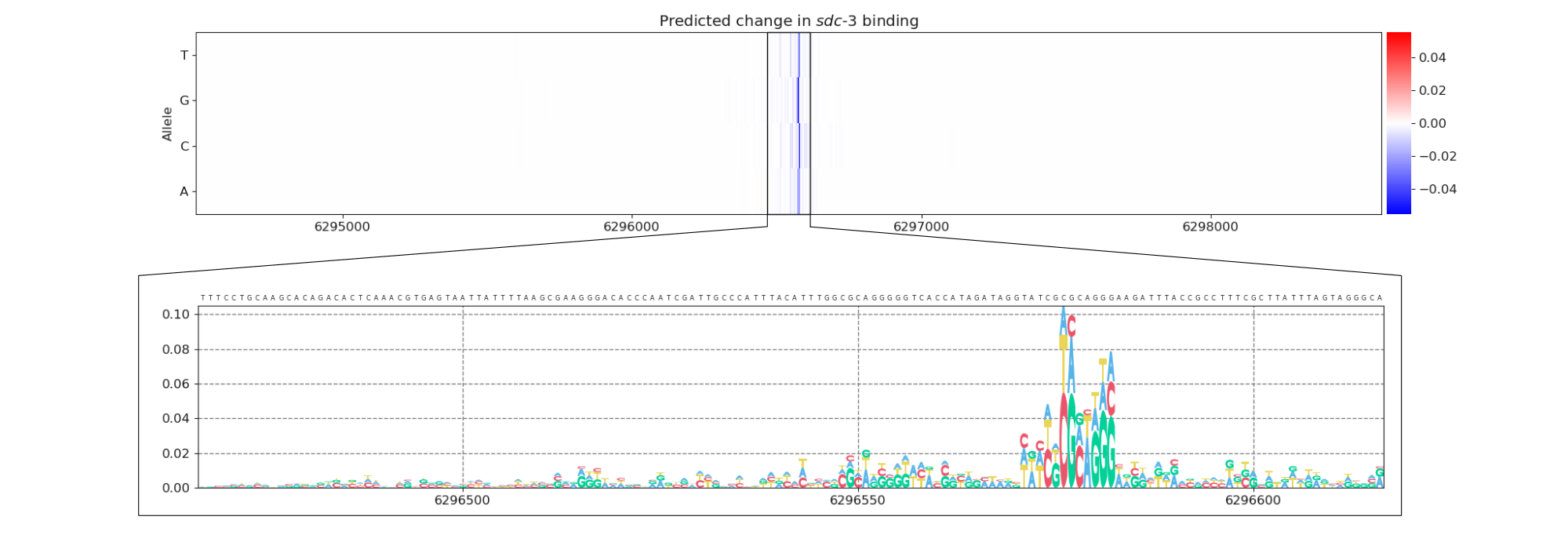

**Supplementary Figure 4: *In silico* saturated mutagenesis applied to the high-confidence DCC binding site in *C. elegans*.** The predicted effect of every possible SNP in the high-confidence DCC binding site (ce11:chrX:6294496-6298590) on the probability of *sdc-3* binding (accession no. SRX059242) relative to the prediction for the reference sequence for all positions in the 4095 bp input sequence (top panel). The same as the top panel, except zoomed in on the center 150 bp, and with the score for a particular allele calculated relative to the lowest probability associated with any allele at its position, so that motif height only measures gain of binding (bottom panel). The canonical TCGCGCAGGG sequence (ce11:chrX:6296574-6296583) appears highly predictive of *sdc-3* binding.

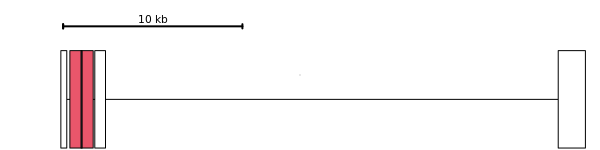

**Supplementary Figure 5:** **Overview of the *T48* gene.** The *T48* gene (dm6:chr3R:26881734-26910997) is shown, and the mesodermal enhancer (dm6:chr3R:26882237-26883537) is drawn as a red box in the first intron. The location of the variants (dm6:chr3R:26882886-26882889) is visualized as a black vertical line occurring near the center of the enhancer.

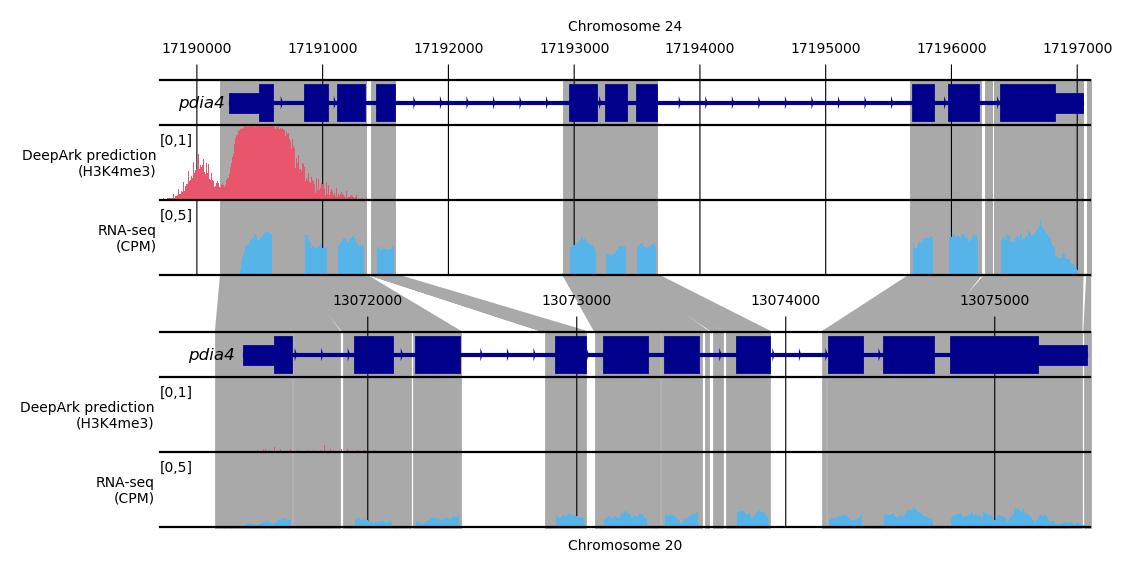
**Supplementary Figure 6: Interspecies predictions with DeepArk indicate diminished *cis*-regulatory activity in *O. latipes* relative to *D. rerio*.** For the *pdia4* gene, which is highly conserved in both *D. rerio* (top) and *O. latipes* (bottom), DeepArk’s feature for H3K4me3 at 6 hpf (accession no. DCD000648SQ) predicts a loss of H3K4me3 at *pdia4*’s promoter in the *O. latipes* genome, which would be associated with diminished or loss of expression at 13 hpf in *O. latipes^32,40^*. Accordingly, normalized coverage counts in RNA-seq from *D. rerio* at 6 hpf and *O. latipes* at 13 hpf shows diminished expression of *pdia4* in *O. latipes* relative to *D. rerio*. CPM, counts per million mapped reads.

**Supplementary Table 1:** All regulatory features predicted with DeepArk, as well as DeepArk’s feature-wise performance on the held-out testing data. Test performance was not calculated for features with fewer than 50 positive examples in the test set.

**Supplementary Table 2:** List of chromosome splits used for training, validation, and testing.

**Supplementary Table 3:** List of variants in the ALDOB enhancer, their reported expression effect in the MPRA, and effects predicted by DeepArk.

**Supplementary Table 4:** The DeepArk features for DCC components used to identify DCC binding on the *C. elegans* X chromosome.

**Supplementary Table 5:** DeepArk’s predictions for each of the T48 enhancer alleles.

**Supplementary Table 6:** The *in vivo* expression effects of the T48 enhancer alleles.

**Supplementary Table 7:** Each *O. latipes* dataset used, along with the accuracy of the matched regulatory features from DeepArk used to perform interspecies regulatory prediction.

**Supplementary Table 8:** The hyperparameters used by each DeepArk model.

| **Species** | **Learning rate** | **Dropout probability** | **Batch size** | **Weight decay** | **Momentum** |
| --- | --- | --- | --- | --- | --- |
| *Caenorhabditis elegans* | 0.1 | 0.15 | 128 | 3.00E-06 | 0.9 |
| *Danio rerio* | 0.1 | 0.2 | 128 | 1.00E-06 | 0.9 |
| *Drosophila melanogaster* | 0.1 | 0.2 | 128 | 3.00E-06 | 0.9 |
| *Mus musculus* | 0.3 | 0.15 | 128 | 1.00E-06 | 0.9 |

**Supplementary Table 9:** Thresholds used to filter datasets for each species.

| **Species** | **Assay Type** | **Target Type** | **Minimum Peaks** | **Minimum Mapped Reads** |
| --- | --- | --- | --- | --- |
| *Caenorhabditis elegans* | DNase-seq | chromatin | 500 | 5000000 |
| *Caenorhabditis elegans* | ChIP-seq | transcription factor | 500 | 2000000 |
| *Caenorhabditis elegans* | ChIP-seq | histone mark – narrow | 500 | 2000000 |
| *Caenorhabditis elegans* | ChIP-seq | histone mark – broad or enriched in repetitive regions | 500 | 5000000 |
| *Danio rerio* | ATAC-seq | chromatin | 2500 | 25000000 |
| *Danio rerio* | ChIP-seq | transcription factor | 2500 | 10000000 |
| *Danio rerio* | ChIP-seq | histone mark – narrow | 2500 | 10000000 |
| *Danio rerio* | ChIP-seq | histone mark – broad or enriched in repetitive regions | 2500 | 25000000 |
| *Drosophila melanogaster* | DNase-seq | chromatin | 500 | 5000000 |
| *Drosophila melanogaster* | ChIP-seq | transcription factor | 500 | 2000000 |
| *Drosophila melanogaster* | ChIP-seq | histone mark – narrow | 500 | 2000000 |
| *Drosophila melanogaster* | ChIP-seq | histone mark – broad or enriched in repetitive regions | 500 | 5000000 |
| *Mus musculus* | DNase-seq | chromatin | 5000 | 50000000 |
| *Mus musculus* | ChIP-seq | transcription factor | 5000 | 20000000 |
| *Mus musculus* | ChIP-seq | histone mark – narrow | 5000 | 20000000 |
| *Mus musculus* | ChIP-seq | histone mark – broad or enriched in repetitive regions | 5000 | 50000000 |

**Supplementary Table 10:** Results from conducting *in silico* saturated mutagenesis for *sdc-3* at the high-confidence DCC binding site on the *C elegans* X chromosome.

**Supplementary Table 11**: Primer sequences used for *T48* enhancer mutants.

| **Name** | **Primer Sequence** |
| --- | --- |
| T48_CAGGAAG_fw | TCGCACGCAGAACTTCCTGCCTCTGGCCATCCC |
| T48_CAGGAAG_rv | GGGATGGCCAGAGGCAGGAAGTTCTGCGTGCGA |
| T48_CAGGTAC_fw | CGCACGCAGAAGTACCTGCCTCTGGCCATCCC |
| T48_CAGGTAC_rv | GGGATGGCCAGAGGCAGGTACTTCTGCGTGCG |
| T48_CAGGCAG_fw | TCGCACGCAGAACTGCCTGCCTCTGGCCATCCC |
| T48_CAGGCAG_rv | GGGATGGCCAGAGGCAGGCAGTTCTGCGTGCGA |
| T48_CAGGTAG_fw | CGCACGCAGAACTACCTGCCTCTGGCCATCCCGCTTGCAC |
| T48_CAGGTAG_rv | GTGCAAGCGGGATGGCCAGAGGCAGGTAGTTCTGCGTGCG |

**Supplementary Table 12:** List of URLs for BAM files from DANIO-CODE.
